# The chaos in calibrating crop models

**DOI:** 10.1101/2020.09.12.294744

**Authors:** Daniel Wallach, Taru Palosuo, Peter Thorburn, Zvi Hochman, Emmanuelle Gourdain, Fety Andrianasolo, Senthold Asseng, Bruno Basso, Samuel Buis, Neil Crout, Camilla Dibari, Benjamin Dumont, Roberto Ferrise, Thomas Gaiser, Cecile Garcia, Sebastian Gayler, Afshin Ghahramani, Santosh Hiremath, Steven Hoek, Heidi Horan, Gerrit Hoogenboom, Mingxia Huang, Mohamed Jabloun, Per-Erik Jansson, Qi Jing, Eric Justes, Kurt Christian Kersebaum, Anne Klosterhalfen, Marie Launay, Elisabet Lewan, Qunying Luo, Bernardo Maestrini, Henrike Mielenz, Marco Moriondo, Hasti Nariman Zadeh, Gloria Padovan, Jørgen Eivind Olesen, Arne Poyda, Eckart Priesack, Johannes Wilhelmus Maria Pullens, Budong Qian, Niels Schütze, Vakhtang Shelia, Amir Souissi, Xenia Specka, Amit Kumar Srivastava, Tommaso Stella, Thilo Streck, Giacomo Trombi, Evelyn Wallor, Jing Wang, Tobias K.D. Weber, Lutz Weihermüller, Allard de Wit, Thomas Wöhling, Liujun Xiao, Chuang Zhao, Yan Zhu, Sabine J. Seidel

**Affiliations:** INRAE, UMR AGIR, Castanet Tolosan, France; Natural Resources Institute Finland (Luke), Helsinki, Finland; CSIRO Agriculture and Food, Brisbane, Queensland, Australia; ARVALIS - Institut du végétal Paris, France; Agricultural and Biological Engineering Department, University of Florida, Gainesville, Florida; Department of Earth and Environmental Sciences, Michigan State University, East Lansing, Michigan; INRAE, UMR 1114 EMMAH, Avignon, France; School of Biosciences, University of Nottingham, Loughborough, UK; Department of Agriculture, Food, Environment and Forestry (DAGRI), University of Florence, Italy; Plant Sciences & TERRA Teaching and Research Centre, Gembloux Agro-Bio Tech, University of Liege, Gembloux, Belgium; Institute of Crop Science and Resource Conservation, University of Bonn, Germany; Institute of Soil Science and Land Evaluation, Biogeophysics, University of Hohenheim, Stuttgart, Germany; Centre for Sustainable Agricultural Systems, Institute for Life Sciences and the Environment, University of Southern Queensland, Toowoomba, Queensland, Australia; Aalto University School of Science, Espoo, Finland; Wageningen University & Research, Wageningen, The Netherlands; Institute for Sustainable Food Systems, University of Florida, Gainesville, Florida; College of Resources and Environmental Sciences, China Agricultural University, Beijing, China; Royal Institute of Technology (KTH), Stockholm, Sweden; Ottawa Research and Development Centre, Agriculture and Agri-Food Canada, Ottawa, Canada; CIRAD, Persyst Department, Montpellier, France; Leibniz Centre for Agricultural Landscape Research, Müncheberg, Germany; Global Change Research Institute CAS, Brno, Czech Republic; Institute of Bio- and Geosciences - IBG-3, Agrosphere, Forschungszentrum Jülich GmbH, Jülich, Germany; INRAE, US 1116 AgroClim, Avignon, France; Department of Soil and Environment, Swedish University of Agricultural Sciences (SLU), Uppsala, Sweden; Hillridge Technology Pty Ltd, Sydney, Australia; Institute for Crop and Soil Science, Federal Research Centre for cultivated Plants, Julius Kühn-Institut (JKI), Braunschweig, Germany; CNR-IBE, Firenze, Italy; Department of Agroecology, Aarhus University, Tjele, Denmark; Grass and Forage Science / Organic Agriculture, Institute of Crop Science and Plant Breeding, Kiel University, Kiel, Germany; Institute of Biochemical Plant Pathology, Helmholtz Zentrum München-German Research Center for Environmental Health, Neuherberg, Germany; Institute of Hydrology and Meteorology, Chair of Hydrology, Technische Universität Dresden, Dresden, Germany; National Institute of Agronomic Research of Tunisia (INRAT), Agronomy Laboratory, University of Carthage, Tunis, Tunisia; National Agronomy Institute of Tunisia (INAT), University of Carthage, Tunis, Tunisia; Lincoln Agritech Ltd., Hamilton, New Zealand; National Engineering and Technology Center for Information Agriculture, Jiangsu Key Laboratory for Information Agriculture, Jiangsu Collaborative Innovation Center for Modern Crop Production, Nanjing Agricultural University, Nanjing, Jiangsu, China

**Keywords:** calibration recommendations, system models, parameter estimation, phenology

## Abstract

Calibration, the estimation of model parameters based on fitting the model to experimental data, is among the first steps in many applications of system models and has an important impact on simulated values. Here we propose and illustrate a novel method of developing guidelines for calibration of system models. Our example is calibration of the phenology component of crop models. The approach is based on a multi-model study, where all teams are provided with the same data and asked to return simulations for the same conditions. All teams are asked to document in detail their calibration approach, including choices with respect to criteria for best parameters, choice of parameters to estimate and software. Based on an analysis of the advantages and disadvantages of the various choices, we propose calibration recommendations that cover a comprehensive list of decisions and that are based on actual practices.

**Highlights:**

- We propose a new approach to deriving calibration recommendations for system models
- Approach is based on analyzing calibration in multi-model simulation exercises
- Resulting recommendations are holistic and anchored in actual practice
- We apply the approach to calibration of crop models used to simulate phenology
- Recommendations concern: objective function, parameters to estimate, software used

## 1 Introduction

Calibration is an important part of the modelling process, since it enables the numerical model results and their reliable use in model applications. It is undertaken in many fields that use system models, including environmental models (Jakeman et al., 2006), hydrological models (Badham et al., 2019), atmospheric models (Steele and Werndl, 2013), models of pest and disease dynamics (Donatelli et al., 2017), and agricultural models (Seidel et al., 2018). Essentially, model calibration adjusts model parameters to reduce the error between the model results and the measured data. The majority of simulation studies involve some type of calibration prior to model application. Calibration is often necessary because parameter values are usually not universally valid, as explained by Fath and Jorgensen (2011) in the context of ecological models, and as explained in the context of crop models, based on statistical principles (Wallach, 2011). Calibration of nonlinear models is a major area of study in statistics (Seber and Wild, 1989; Sen and Srivastava, 1990), but system models have several features which make calibration particularly challenging (Wallach et al., 2019). Firstly, system models often have a large number of parameters, often many more than the number of observed data, which means calibration of all parameters is not possible. So one must decide which parameters to estimate (Doherty and Hunt, 2009; Necpálová et al., 2015). Even when one estimates a subset of the model parameters, there is often a problem of equifinality, meaning that various different combinations of parameter values can give the same results, and so calibration does not lead to unique parameter values (Beven and Freer, 2001). Furthermore, system models usually simulate multiple different variables which can be compared with observed data, leading to the problem of combining information about the fit of the different variables into a single criterion for calibration, or possibly of defining multiple criteria (Wöhling et al., 2013b). Software used for the calibration is an additional problem. Often one ‘externally’ couples the existing model software to calibration software (Buis et al., 2015; He et al., 2010; Hunt et al., 1993), but this can require substantial effort. As a result, calibration for crop models is often done by manual trial and error without using an automated routine (Seidel et al., 2018), though this could also be a deliberate choice to exert more supervision over the calibration.

In response to these and other difficulties, there have been numerous studies published concerning calibration recommendations for system models in multiple disciplines. One type of study focuses on a particular model; it identifies the most important parameters in that model, and explains how they can be estimated from data (Ahuja et al., 2011). Other studies have focused on the implementation of a Bayesian approach or on the comparison of frequentist and Bayesian approaches (Gao et al., 2020; Jansen and Hagenaars, 2004; Sexton et al., 2016; Van Oijen et al., 2005), on numerical methods of seeking best parameter values (Bhar et al., 2020; Franchini and Galeati, 1997; Madsen et al., 2002), on the choice of parameters to estimate (Angulo et al., 2013), on the definition of multiple objective functions as a way of handling multiple simulated responses (Efstratiadis and Koutsoyiannis, 2010) or on which observed data to use for calibration (Hunt et al. 2001). There do not seem to be calibration recommendations based on a holistic view of model calibration for system models, such that the recommendations cover the full range of decisions that calibration involves, and that are based on actual practice. A holistic treatment is of importance, because potentially any or all of the decisions involved in the calibration process might have an important impact on the results. It is important to base recommendations on the range of actual practices, to ensure that a wide range of feasible approaches is considered.

The specific system models considered here are crop models, which consist of a set of mathematical equations representing physically based or (semi)empirical processes that describe plant development and growth as well as soil conditions as affected by weather, soil characteristics and crop management. Crop models are widely used to study, understand, and optimize crop production in current and future environments (Ewert et al., 2015; Keating and Thorburn, 2018; Tsuji et al., 1998). In our study, we focus specifically on the use of crop models to simulate crop phenology i.e. the cycle of biological events in plants, because matching the phenology of crop varieties to the climate in which they grow is a critical crop production strategy (Hunt et al., 2019; Rezaei et al., 2018, 2015). In general, the simulation of crop phenology is an essential part of crop models and implemented as a phenological model component (or submodel) in the crop model. In many crop modelling studies, the focus is specifically on simulating phenology (Gao et al., 2020; Kawakita et al., 2020; Wu et al., 2017) but also calibration of crop models using just phenological observations can be a first step in crop simulation studies (Kimball et al., 2019; Li et al., 2015). Modeling plant phenology is also important in understanding ecosystem response to global warming (Piao et al., 2019)

The objective of the present study is to define a novel approach to developing recommendations for calibration of system models, and to apply it to derive recommendations for calibration of phenology simulation using crop models. The approach considers the full range of decisions involved in the calibration process and is based on information about the actual choices made by multiple modeling groups. The proposed approach involves three steps. First, one or more multi-model simulation studies are organized, where all participating modeling teams are given the same data for calibration and asked to provide simulated values for the same outputs using their usual calibration technique. Secondly, each team is asked to complete a detailed questionnaire about the choices made for each calibration decision. This provides information about the range of choices that are made in practice. Finally, the advantages and disadvantages of the different choices for each calibration decision are analyzed, which provides the basis for a set of recommended practices.

## 2 Materials and Methods

This study is based on two multi-model simulation studies. The studies were organized within the framework of the Agricultural Modeling Intercomparison and Improvement Project (AgMIP, Rosenzweig et al.; 2013). The co-leaders of the calibration activity designed the studies as a way of obtaining information on crop model calibration practices, when the objective is simulation of crop phenology. They invited crop modeling groups to participate by announcements on the AgMIP website and in messages to the mailing lists of several widely used models. All modeling groups that asked to participate were accepted.

We use here the term “model structure” to designate a specific set of model equations. The model structures used by the participants are listed in Supplementary Table S1. We speak of modeling group to designate the group of researchers that implemented the model structure in a specific case. The modeling group was responsible for determining all aspects of the calibration procedure and also for choosing the values of fixed parameters, i.e. those not determined by calibration. Twenty-seven modeling groups participated in the first simulation study, based on data from wheat fields in France. Twenty-six of those groups, and two additional groups, participated in the second simulation study, based on data from wheat fields in Australia. Thus overall 29 different groups participated in at least one of the studies.

The modeling groups are identified as M1-M29 and the same identifier is used for the same modeling group working with the French and Australian data. The name of the model structure used by each group is not given, since this might give the erroneous impression that the calibration approach and simulation results are specific to that model structure, while in fact they depend on both the model structure and the decisions made by the modeling group. Three of the model structures, coded as S1, S2, and S3, were used by multiple groups. Structure S1 was used by four (French dataset) or three (Australian dataset) groups, structure S2 by three groups and structure S3 by two groups. Comparisons within these three groups provide information about the variability as to calibration approach between different modeling groups calibrating the same model structure.

Figure 1 shows schematically the flow of information in the multi-model exercises. In each study, participants were given the input data usually required for running a crop model, namely daily weather data, information on crop management, and information on soil characteristics, for multiple environments. An “environment” refers to a specific combination of site and sowing date. The modeling teams were also given data on wheat phenology from a subset of those environments for calibration (the “calibration” environments) and asked to provide simulated phenology for the remaining environments, the “evaluation” environments.

**Figure 1.**
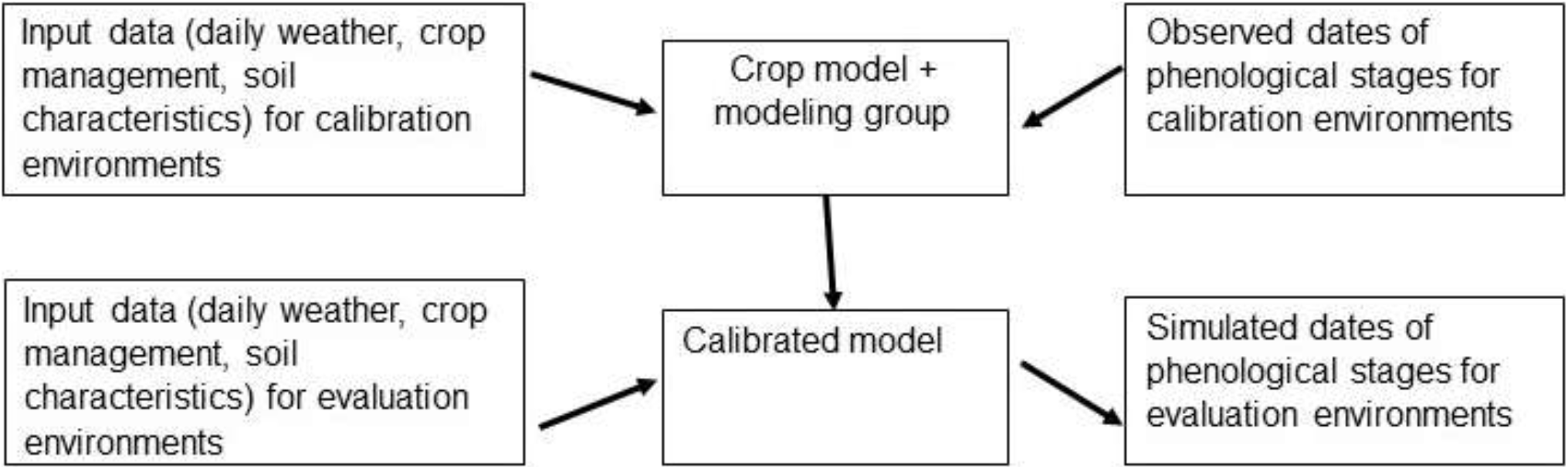
Schematic diagram of the two multi-model simulation studies.

The observations in the French dataset were for days from sowing to two phenological stages, namely beginning of stem elongation (growth stage 30 on the BBCH and Zadoks scales) and middle of heading (growth stage 55 on the BBCH and Zadoks scales) (Table 1). These two stages are of practical importance because they can easily be determined visually and are closely related to the recommended dates for the second and third N fertilizer applications in France. Observed data were provided for two varieties, namely Apache and Bermude. In all cases, the modeling groups used the same calibration approach for both varieties. Therefore, in this study we only report a single calibration approach for each modeling group for the French dataset. The modeling groups were asked to provide simulated values for those same two growth stages for the evaluation environments.

**Table 1.** Description of datasets. The French dataset was repeated for two varieties (Apache and Bermude). The numbers shown represent the days from sowing to the indicated stages on BBCH or Zadoks scale. An environment corresponds to a specific combination of site and sowing date.

| Dataset | Calibration data |  | Evaluation data |  |
| --- | --- | --- | --- | --- |
|  | Number of environments | Observed or interpolated phenological stages | Number of environments | Observed or interpolated phenological stages |
| French<br>(repeated for varieties Apache and Bermude) | 14 | BBCH30, BBCH55 | 8 | BBCH30, BBCH55 |
| Australian<br>(variety Janz) | 24 | Each integral Zadoks stage from first to last observed Zadoks stage | 18 | Z30, Z65, Z90 |

The Australian dataset resulted from measurements for the Zadoks growth stage (Zadoks et al., 1974), that were recorded every two weeks in each environment. The data were interpolated, to give days from sowing to every integer Zadoks stage from the first to the last observed stage, which were the data provided to each modeling group for the calibration environments. For the evaluation environments, participants were asked to provide the simulated values for the number of days from sowing to stages Z30 (Zadoks stage 30, pseudostem, i.e. youngest leaf sheath erection), Z65 (Zadoks stage 65, anthesis half-way, i.e. anthers occurring half way to tip and base of ear), and Z90 (Zadoks stage 90, grain hard, difficult to divide). These stages are often used for management decisions or to characterize phenology.

At no point did the participants have access to the evaluation data. Furthermore, none of the sites or years represented in the evaluation environments occurred in the calibration environments. Thus this was a rigorous evaluation of how well the modeling groups were able to simulate wheat phenology for new environments. Details for the evaluation results are provided in (Wallach et al. (2021b, 2021a). The prediction errors are also summarized in Supplementary Table S7.

In both simulation exercises, each participating modeling group was asked to calibrate the model in their “usual” way, using the calibration data provided. Each group was also asked to complete a questionnaire, detailing how the calibration was conducted (Table 2). The questions were chosen to cover as completely as possible the full set of the decisions that are associated with the calibration approach, and to provide information about the underlying reasoning for the choice of modeling approach (see questions 3 and 6).

**Table 2.**
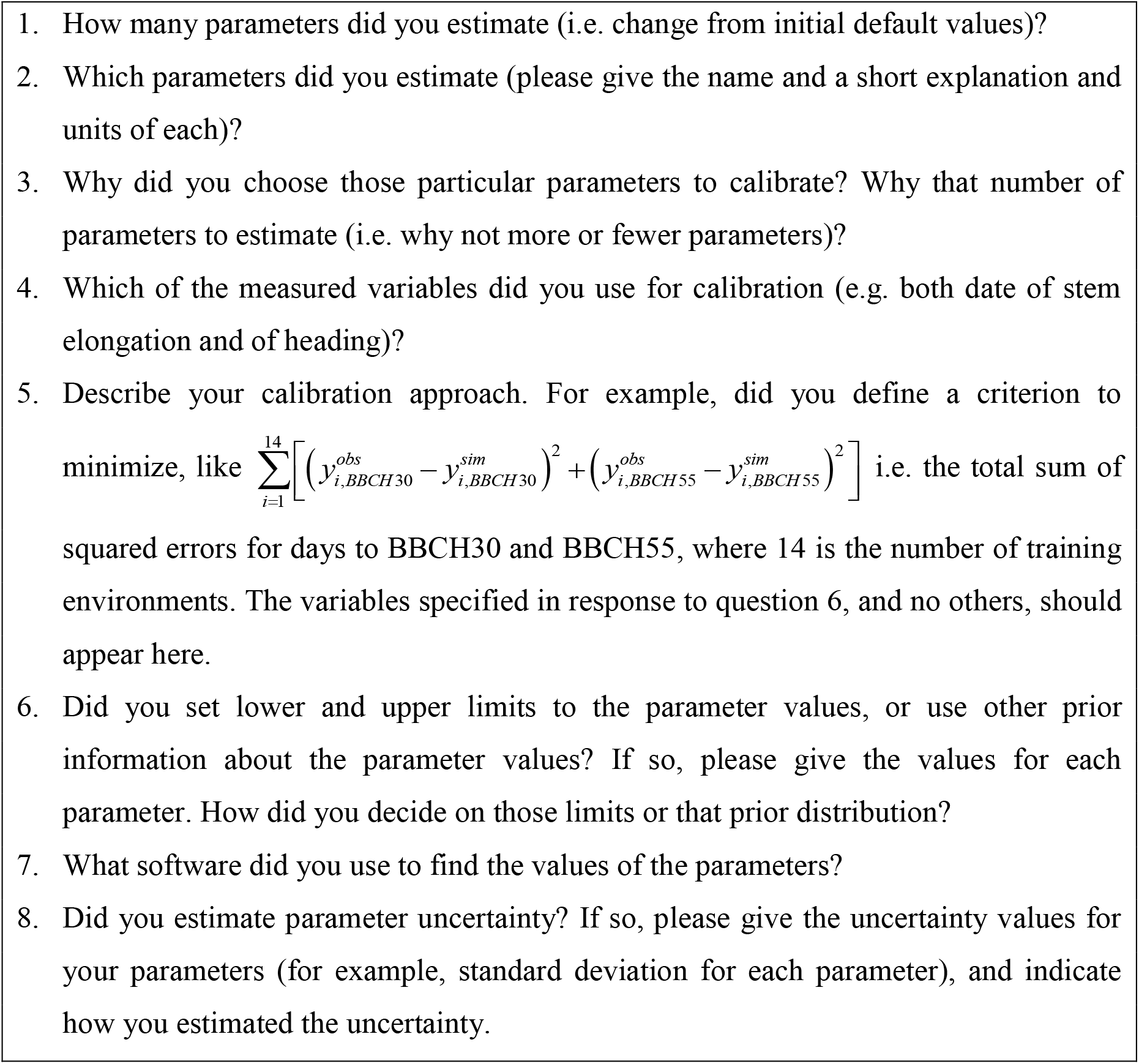
The questions related to calibration approach in the questionnaire filled out by all participating groups. The full questionnaire is presented in Supplementary Table S2.

For details on how each model structure simulates phenology, see the references for each structure in Supplementary Table S1. Here, we provide only a short overview, with emphasis on the parameters involved. The basic output of phenology simulation is days from sowing to various phenological stages or between various phenological stages. The number and identity of simulated stages varies with the model, and may include both observable stages (for example anthesis) and stages that are model constructs (for example, start of linear phase of grain filling). The most important inputs that determine spring wheat phenology are daily temperature and photoperiod (Aslam et al., 2017), while for winter wheat it is also important to include the process of vernalization, i.e. the effect of low winter temperatures on development (Li et al., 2013). Most model structures take into account all three factors, though not all factors affect the development rate at all stages of development. Some structures take into account only temperature, or temperature and either photoperiod or vernalization. Most model structures take temperature into account by calculating thermal time, calculated most simply as the sum of the daily temperature above some threshold temperature (a parameter). In other models the daily contribution to degree days may have a plateau above some optimal temperature (a parameter), or decline above the optimum temperature at some rate (a parameter) or be some more complex function of temperature (Kumudini et al., 2014; Wang et al., 2017). The parameters of the temperature response curve may differ at different stages in the development cycle. Wheat is a long-day plant, which flowers earlier in longer days. Often phenology response to photoperiod is modeled using two parameters, a threshold photoperiod below which development rate increases with increasing photoperiod and a sensitivity coefficient, which describes the rate of increase, though other functions, with other parameterizations, are also used. Vernalization is described as a period of low temperatures, which must be experienced before the plant can flower. Vernalization parameters can include the upper limit for temperature to count as vernalizing, and the required number of vernalizing days. To determine the day of occurrence of a given stage, most models have an internal counter of physiological time, which often is based on degree days modulated by photoperiod. When this counter attains some predetermined value specific to the stage in question, which is a model parameter, that stage is deemed to have been attained. Some models also relate development to the rate of leaf appearance (called the phyllochron, a parameter) or rate of tillering. Several models account for cold or drought stress in the simulation of the development rate. If the development rate depends on drought stress, then it is sensitive to the parameters in the model that determine soil water content and soil-plant water dynamics. In other models, phenology is independent of the model parameters that determine growth and soil dynamics.

## 3 Results

The decisions required for calibration can be divided into three groups. I) decisions related to the criterion that defines the best parameter values, II) decisions related to the choice of parameters to be estimated, and III) decisions related to the numerical calculation of the best parameter values. The more detailed decisions within each group are shown in Figures 2–4, which also indicate the choices made by the participating modeling groups and the number of groups that made each particular choice. Details for each individual modeling group are shown in Supplementary Tables S3, S4, and S5 for choices concerning the criterion of best parameter values, the parameters to estimate, and the algorithm and software, respectively.

**Figure 2.**
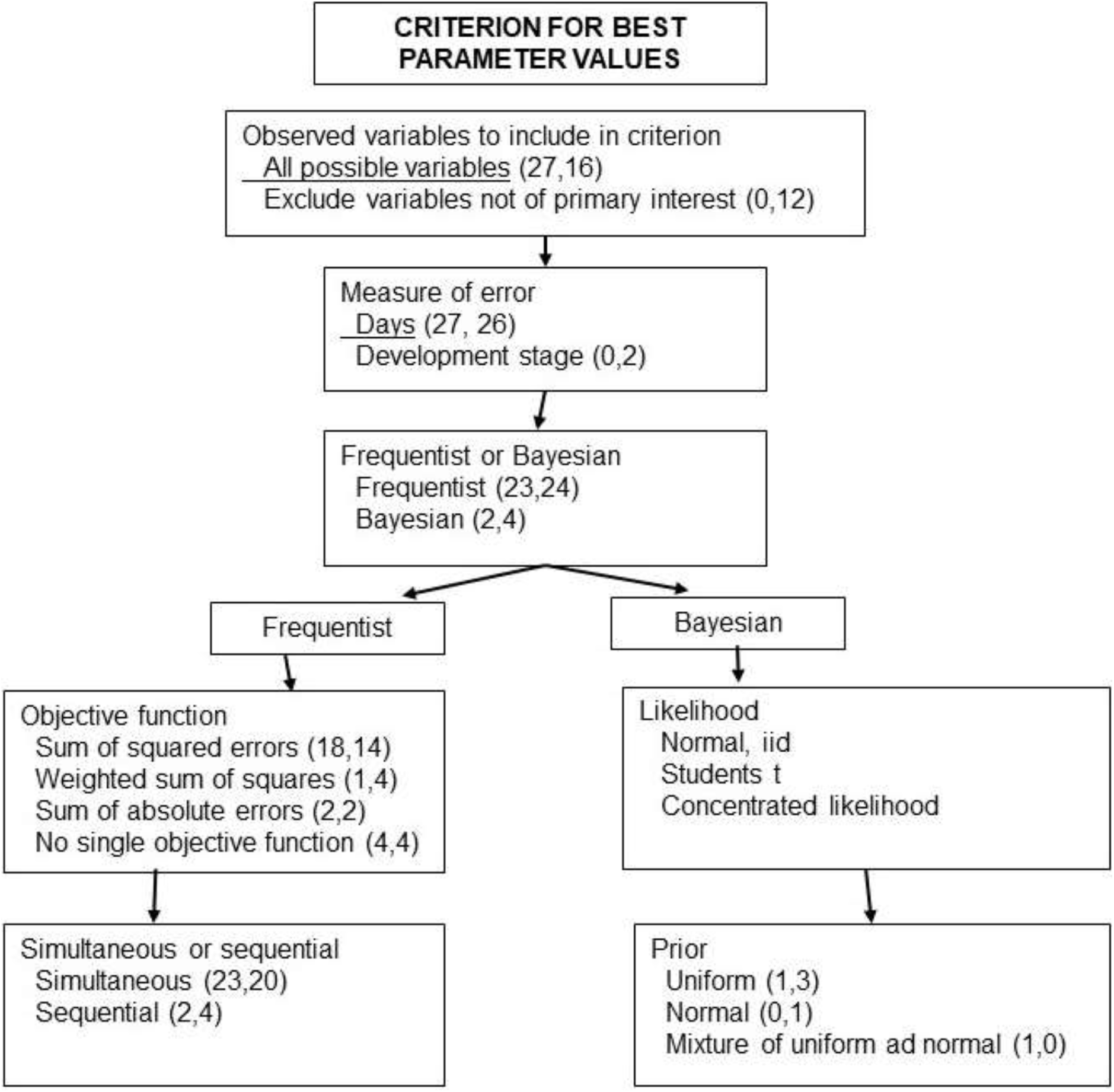
The calibration decisions related to the criterion for best parameter values, and the choices made by multiple modeling groups in two studies. The numbers in parentheses are the numbers of groups that made the indicated choice for the French and Australian data sets, respectively. Choices recommended here are underlined.

### 3.1 Criterion for best parameters

A first calibration decision in this category is which variables to use in the criterion that defines the best parameter values, and in particular whether to use only those variables for which predictions are sought, or also additional observed variables. By variable, we mean the type of data. For example, days to flowering and days to heading would be two different variables. The French dataset only had observations for two variables, namely days to phenological stages BBCH30 and BBCH55 (see Table 1), and groups were requested to report the simulated values for those same two variables, for both the calibration and evaluation environments. Thus, it was a logical consequence that for almost all groups the criterion of best parameters included observations of both those variables. Two groups (M9 and M18) used model structures that did not simulate the number of days to stage BBCH30, so these groups only used a subset of the observed variables (i.e. the observations of days to stage BBCH55) in the criterion defining best parameters. The Australian dataset, on the other hand, had many observed variables (i.e., days to many phenology stages), while evaluation was based just on simulated days to three stages (Z30, Z65, and Z90). Here, the choice of variables to include in the criterion was not so straightforward. For the Australian dataset about 40% of the groups used only the variables to be simulated (days to Z30, Z65, and Z90 or a subset if the model structure did not simulate all those variables), while about 60% of the modeling groups included other observed variables in the criterion of best parameters. Different groups that used the same model structure did not necessarily make the same choice here. Considering structure S1, used by groups M2, M3, and M4 and the Australian dataset, all three groups used minimum sum of squared errors as the criterion defining the best parameter values. However, group M2 included additional variables in addition to the variables to be simulated in their sum of squared errors. Group M3 used only squared errors for Z65 and Z90, and group M4 used squared errors for Z30, Z65, and Z90.

A second decision concerns the definition of error. Almost all groups expressed error in terms of days to reach a specified stage (Table S3). However, it is also possible to express error in terms of phenological stage. In the simplest case, suppose that a model structure calculates Zadoks stage each day (i.e., the internal counter each day is directly or is translated into a value for Zadoks stage). Suppose that for a particular environment it is observed that stage Z30 is attained on day 45, but the simulated day is 40. The error in days is 5 days. Suppose that the simulated stage on day 45 is Z33.4. Then the error in terms of development stage is 30-33.4=−3.4. For the French dataset all groups calculated error in days, but for the Australian dataset three groups expressed error in terms of development stage rather than days.

A third decision is whether to use a frequentist or Bayesian perspective. If a frequentist perspective is chosen, one must define the mathematical form of the objective function. F a Bayesian perspective is chosen, one must define the form of the likelihood and the prior distributions for the parameters. The large majority of groups followed a frequentist approach, where the estimated parameter values are those values that minimize some measure of error between the simulated and observed values (Table S3). Most of the frequentist groups sought to minimize the sum of squared errors, where the sum is over calibration environments and over all variables included in the criterion. This is the ordinary least squares (OLS) criterion. One (French data) or four (Australian data) groups used a different measure of distance between observed and simulated values, namely the sum of root mean squared errors for the different variables, or a weighted sum of squared errors. Two groups chose to minimize the sum of absolute errors. This is the least absolute value criterion (LAV). Four groups for the French dataset, and the same four groups for the Australian dataset, did not define an explicit objective function to be minimized, but rather sought parameter values to give a “best fit” to the data, where “best fit” was determined visually or by some subjective combination of mean squared error, R^2^, or other fit metrics. Groups using the same model structure did not necessarily make the same decisions. For example, among the three groups that used model structure S2, for the Australian dataset, one used the OLS criterion and two had no explicit objective function.

Another decision for the frequentist perspective is whether to fit all the observed variables in a single calculation step or to use multiple steps, adjusting parameters to different variables in each step. Almost all groups estimated all parameters simultaneously (Table S3). However, two (French data) or four (Australian data) groups estimated parameters in more than one step, fitting for example three parameters to the BBCH30 data in the French dataset, and then fixing those parameters at their estimated values and fitting another parameter to the BBCH55 data. Again, groups using the same model structure did not always make the same decisions. For example, among the two groups that used model structure S3, for the Australian dataset, group M23 estimated all parameters simultaneously, while group M24 estimated parameters in two steps.

Of the total of six Bayesian calibrations (two for the French dataset, four for the Australian dataset), three assumed a normal distribution of errors and one a Student’s *t* distribution. One group worked with the concentrated likelihood, which replaces the model variance for each variable by its maximum likelihood value. For the Bayesian groups, parameters were assumed to have either uniform or truncated normal prior distributions.

No group took correlations of errors for different variables in the same environment into account. That is, all groups treated all the errors as though they were independent. Only two groups (M19, M21, see Table S3) took into account the possibility of different error variances for different variables, M21 by using the method of concentrated likelihood (Seber and Wild, 1989) and M19 by dividing the likelihood for each variable by the number of observations of that variable.

### 3.2 Choice of parameters to estimate

Each model structure is parameterized differently, so it is not possible to directly compare names of parameters between model structures. It is, however, possible to identify the role of estimated parameters in the model and base the comparison between groups on that. Details related to the choice of parameters by each group are given in Supplementary Table S4.

Most groups estimated at least some parameters that concern the physiological time required to attain one or more phenological stages. Fifteen groups for both datasets estimated one or more parameters related to vernalization, and 13 groups for both datasets estimated one or more parameters related to photoperiod sensitivity. A smaller number of groups estimated parameters related to the temperature response function (for example minimum temperature or optimum temperature for development) or to tillering or leaf appearance rate (phyllochron). Two groups for each dataset estimated parameters related to the effect of stress on the development rate, and two (French dataset) or three (Australian dataset) groups estimated parameters related to time to emergence.

The number of estimated parameters ranged from one to nine for the French dataset and from two to ten for the Australian dataset. In most cases, the choice of parameters to estimate was based on expert opinion, but four groups for each dataset combined expert opinion with data-based information (for example, testing various combinations of parameters to see which gives the best fit). Five (French dataset) or four (Australian dataset) groups based the choice of parameters to estimate on sensitivity analysis. We have identified the use of expert knowledge and data-based choice of parameters to estimate as two separate categories, but it should be noted that expert knowledge by itself usually adapts the choice of parameters to estimate to the data, at least to some extent. This can be seen from the fact that almost every group that based the choice of parameters on expert knowledge estimated a different (and in most cases larger) set of parameters based on the Australian dataset, with more observed variables, than based on the French dataset (Figure 5).

Unlike the specific choices of parameters to estimate, the underlying basis for choosing parameters to estimate is applicable in general for system models. It is thus these choices that are shown in Figure 3. There were important differences even between groups using the same model structure. Consider for example structure S2. The three groups that used this model structure (M7, M12, and M13) estimated respectively four, three, and two parameters for the French dataset and nine, four, and two parameters for the Australian dataset. Two of those groups based the choice on expert opinion, while group M7 made a partially data-driven choice.

**Figure 3.**
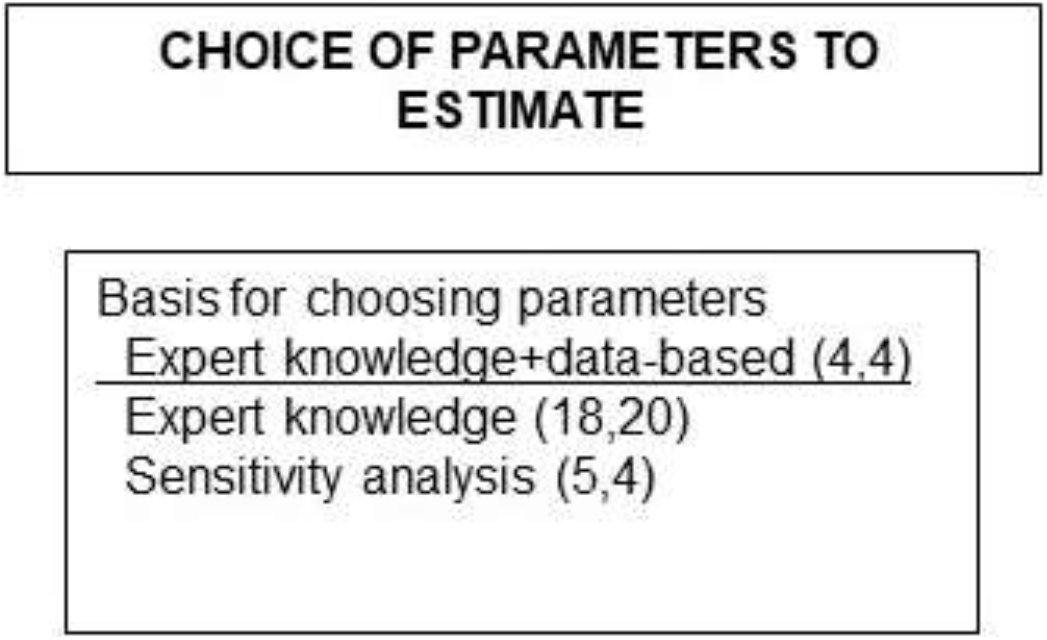
Possible choices of underlying basis for choosing the parameters to estimate, and the choices made by multiple modeling groups in two studies. The numbers in parentheses are the numbers of groups that made the indicated choice for the French and Australian data sets, respectively. The choice recommended here is underlined.

### 3.3 Numerical methods

The basic decision here is the algorithm to use for estimating the parameters (Figure 4). A second, practical decision is the software to use to implement that algorithm. The choices made by each modeling group are shown in Supplementary Table S5, and information about the specific software used is given in Supplementary Table S6. Among the groups that chose a frequentist approach, slightly over half used trial and error to search for the optimal parameter values. In some of those cases, available software was used as an aid, but the final values were found by simply trying different parameter values. The remaining half was split between groups that used a derivative-free search algorithm, usually an algorithm designed to find a global optimum, and those that used a gradient-based algorithm. Many different software solutions were used, including multi-purpose software packages as well as software written expressly for calibration of that particular model structure. Four groups (French data) or six groups (Australian data) used a Markov-Chain Monte-Carlo (MCMC) algorithm to estimate the posterior distribution, using various software packages. That included one group that used an MCMC algorithm though the objective was to minimize the sum of absolute errors. The groups that used the same model structure, in general, did not use the same algorithm and software. For example, considering the two groups using model structure S3, group M23 used a combination of global and local search algorithms and available software packages, while group M24 used trial and error and no software packages.

**Figure 4.**
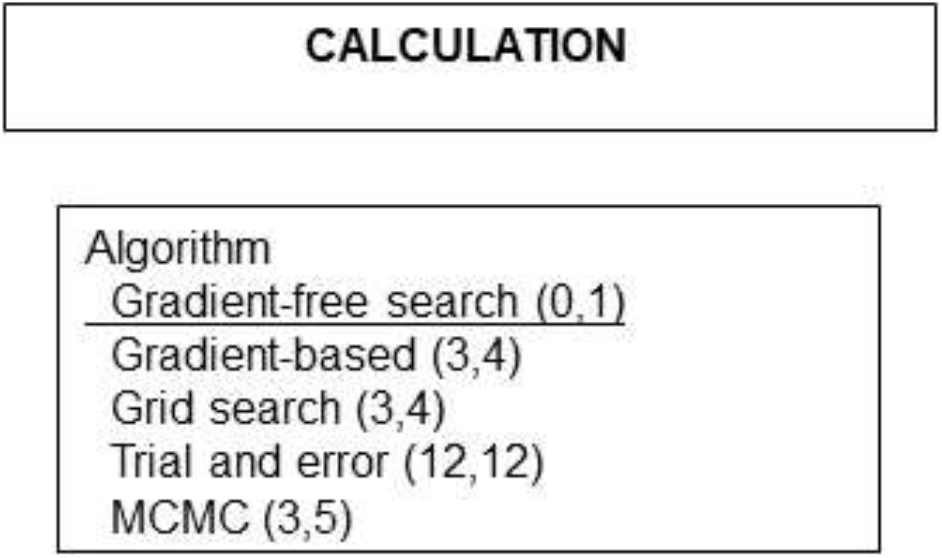
Algorithm type for calculating the parameters, and the choices made by multiple modeling groups in two studies. The numbers in parentheses are the numbers of groups that made the indicated choice for the French and Australian data sets, respectively. Choices recommended here are underlined. For software choices, see Supplementary table S6.

**Figure 5.**
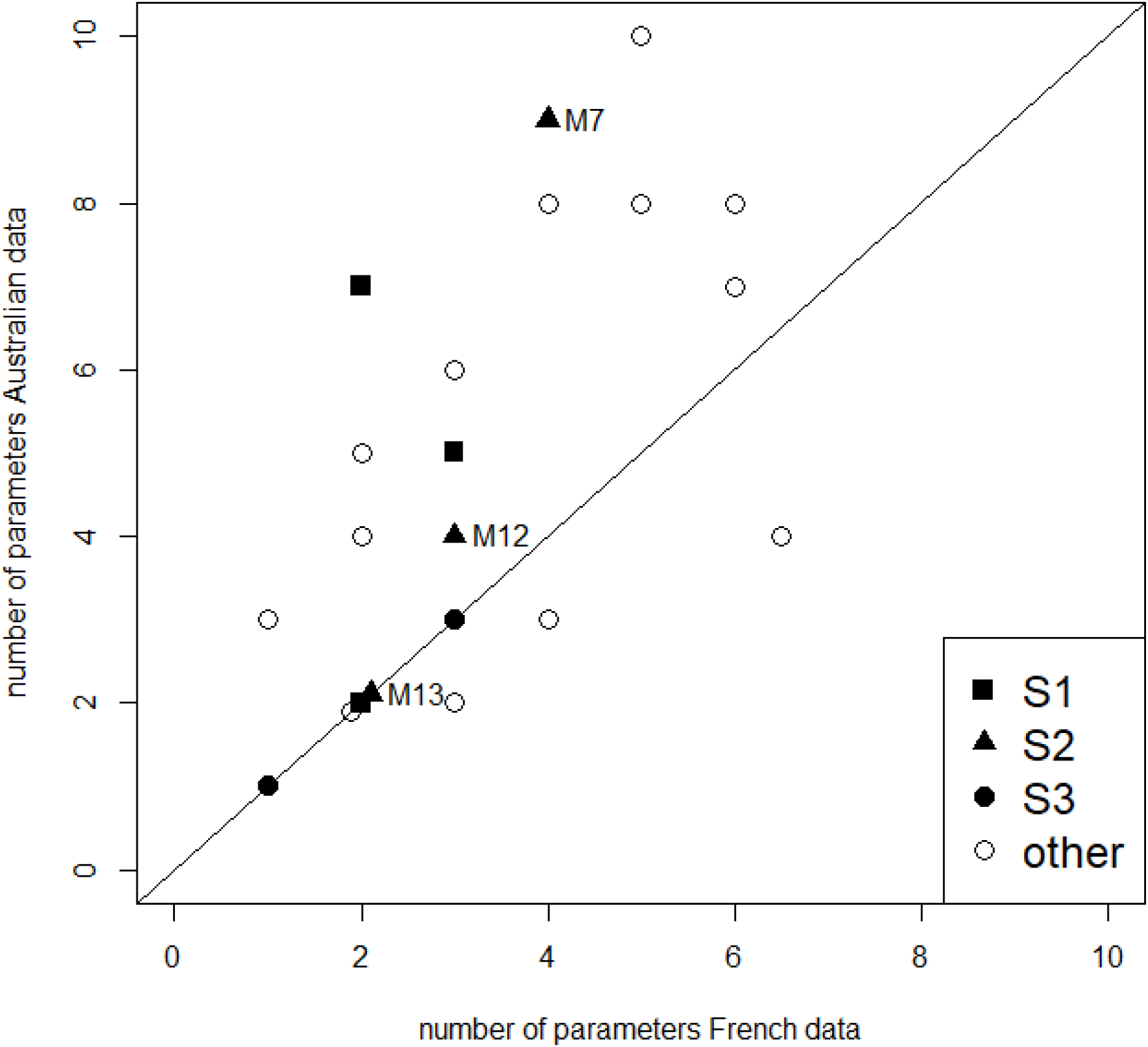
Number of parameters estimated by each modeling group, for the French and Australian datasets. The modeling groups that used model structures S1-S3 are identified.

### 3.4 Level of error

Supplementary Table S6 shows mean absolute error (MAE), where the average is over the calibration data or over the evaluation data, for each modeling group, for the French and Australian datasets. The error results have previously been published (Wallach et al., 2021b, 2020). MAE can be quite different, even for different groups using the same model structure. For example, four modeling groups used model structure S1 for the French dataset, and had MAE for the calibration data of 3.9, 4.9, 6.8 and 12.8 days, respectively.

## 4 Discussion

There is substantial variability in calibration approach between modeling groups, even between groups that use the same model structure. Thus, a first overall conclusion is that we are far from having a consensus on how to calibrate crop models, even for a given model structure and dataset, and even for the relatively simple case which focuses just on phenology. In the following, we discuss the advantages and drawbacks of each choice for each calibration decision, and on that basis make recommendations for good practices. We do not base these recommendations on the levels of error of the different modeling groups, because there is no simple relation between calibration approach and resulting error. Differences in error could be due to differences in model structure, to differences in parameter values for those parameters not estimated by calibration, as well as to differences in any or all of the calibration decisions made by the individual groups.

### 4.1 Criteria for best parameters

A major calibration decision is the list of observed variables (here observed development stages) to include in the criterion of best fit. From a modeling point of view, using as many variables as possible for fitting the model reduces the risk of “getting the right answer for the wrong reason”, i.e. getting a good fit for some variables while other variables, that describe other aspects of system behavior, are poorly simulated (Meyer Oliveira et al., 2021; Wang et al., 2011). Fitting the model to more variables will reduce the aspects of the system that could unknowingly be poorly simulated. This is important, since the same calibrated model might be used for more than one specific purpose. More generally, process-based crop models are argued to be meaningful tools for understanding crop growth and production in response to climate variability and change (Keating and Thorburn, 2018), particularly as they cover interconnections of different system variables in their structures (Ewert et al. 2015). Calibration using multiple observed variables should improve the representation of these interconnections. From a statistical point of view, more data in general leads to predictors with smaller variance, which argues for using all the available data. However, this assumes that the model is correctly specified in the statistical sense, meaning that model errors have expectation zero for all values of the explanatory variables. It has been argued that crop models are most likely statistically incorrectly specified, and as a result, the best parameters for predicting one variable may be different than the best parameters for predicting a different variable (Wallach, 2011). In that case, using additional variables in the objective function may degrade predictive accuracy for the variables of primary interest. This was found to be the case in the study of Guillaume et al. (2011). In fact, it has been suggested that the differences in calibration results using different observed variables could be a diagnostic tool for model structural errors (Wöhling et al., 2013a). If, however, one is willing to assume that statistical misspecification is not too extreme, then it seems worthwhile to include as many of the observed variables as possible in the objective function, so here this is the recommended practice. This does not imply that all variables should have equal weight in the criterion for best parameters. Statistical theory shows that if different variables have different error variances, then that should be taken into account.

Most groups defined error as the difference between the simulated and observed days to reach a given phenological stage, but in a few cases error was defined as the difference between the simulated development stage and the observed stage, on the day of observation. This option requires that the model include some internal counter whose observed and simulated values at observed phenological stages are known, but this is often the case. It has been argued that the problem of minimizing errors is much better behaved numerically when errors are in terms of development stage rather than days (Wallach et al., 2018). On the other hand, one is usually interested in how large the error is in days so this is a more intuitive error measure. Furthermore, this does not require that the model have an internal measurement of development stage. We suggest then to measure error in days.

For the groups that adopted a frequentist perspective, the large majority framed the problem as an ordinary least squares (OLS) problem. Two modeling groups chose parameters by minimizing the sum of absolute errors, which has been argued to have advantages over OLS, as it is less sensitive to outliers (Willmott and Matsuura, 2005). However, if evaluation is based on squared error, then OLS is the more logical choice and is recommended. Four groups did not have an explicit objective function. One obvious disadvantage of this approach is its subjectivity, adding uncertainty in the definition of best-fit to other uncertainties in calibration. A second disadvantage is that one cannot automate the search for the best parameters.

In most cases, a single objective function, combining all errors, was used. In a few cases, however, parameters were fitted sequentially (first to one variable then to the next etc.). This sequential technique has often been recommended for full crop models (L.R. Ahuja et al., 2011; Anothai et al., 2008). This simplifies the mechanics of finding the best parameter values, but it will lead to sub-optimal results with respect to an overall objective function. If the objective is to minimize the total sum of squared errors, for example, the best parameter values are those that minimize exactly that objective function. Necpálová et al. (2015) similarly recommended simultaneous estimation even for multiple observed variables, for an ecosystem biogeochemical model.

A few groups chose a Bayesian rather than a frequentist perspective. There are fundamental differences between frequentist and Bayesian approaches (Berger and Bayarri, 2004). However, for the practical prediction problem here, there are also important similarities. A major difference is that the Bayesian approach focuses on the posterior distribution, which is a distribution of predicted values, while the frequentist approach focuses on point predictions, i.e. one single predicted value. Here, however, all groups were asked for point predictions, so the groups that used Bayesian approach had to choose a single result from the posterior distribution. In all cases, they chose the parameter values that maximized the posterior distribution, which then plays the same role as the objective function for the frequentist approach. Another important difference is that for the Bayesian approach the prior information about parameter values is included in the calculation, while this is not in general included in a frequentist approach. However, in almost all cases here, the frequentist approach included lower and upper bounds on the parameter values (table S5), which is also based on prior information. In fact, a Bayesian approach with normal likelihood and uniform priors leads to exactly the same criterion for best fit, namely minimum squared error subject to the constraints on the parameters, as OLS with bounds on the parameter values.

If the main objective is to obtain a point predictor, then we suggest using a frequentist approach, since adopting a Bayesian approach and calculating a posterior distribution may require unnecessarily long calculation times. If one is also interested in uncertainty information, as it is often the case, then a Bayesian approach has advantages as far as parameter uncertainty is concerned, since the posterior distribution is directly a representation of the uncertainty in the parameter vector. However, it is important to keep in mind that parameter uncertainty is only part of overall uncertainty, and in fact it has been found in several cases that parameter uncertainty is quite a bit less than structure uncertainty (Zhang et al., 2017).

In almost all cases, the calibration approach was directly based on regression methods in statistics, either frequentist or Bayesian. This seems logical, insofar as these statistical methods have desirable properties. However, these properties in general require that certain assumptions be satisfied. The standard assumptions for the OLS method are that the model errors be independent and identically distributed, with expectation 0 (Seber and Wild, 1989; Sen and Srivastava, 1990). For the Bayesian methods, one must make explicit assumptions about the distribution of errors, including whether all errors have the same distribution and whether errors for different variables are correlated. In the case of crop models, with multiple observed variables in each environment, the assumptions of independent identically distributed errors with expectation 0 are not likely to be satisfied (Wallach et al., 2019). Most obviously, errors for different variables in the same environment (e.g., days to development stages Z30 and Z65) may be correlated, since any particularities of the environment affect all variables for that environment. Furthermore, the errors for different variables may have different variances, which violates the assumption that all errors have the same distribution. No group took correlations of errors into account and only two groups took into account the possibility of different variances for errors of different variables.

In general, it would be worthwhile to go a step further in applying statistical methods, beyond employing standard techniques, in order to examine whether the standard assumptions about model error are satisfied. To detect unacceptably large violations of the standard assumptions, one should examine the model residuals (observed minus simulated values) after calibration, as is standard procedure in regression (see for example NIST/SEMATECH, 2013). One should examine overall bias of model residuals, which should be zero, the variances of residuals for different variables, which should be similar, and correlations between residuals for different variables in the same environment, which should be small.

Only phenology data were available in the datasets here, and thus all errors had the same units (days or phenological stage). In cases where variables with different units are observed, for example days to phenological stages and yield, it is meaningless to simply combine errors. In that case, a first step could be to divide all simulated and observed values by an estimated standard deviation of error for that variable, as in weighted least squares (Seber and Wild, 1989). Then all errors would be unitless and could be combined. However, it would still be important to test residuals after calibration.

### 4.2 Choice of parameters to estimate

Of particular interest is the rationale behind the choice of parameters to estimate, and what this implies for the adaptation of the choice of parameters to the dataset. In most cases, the choice of parameters to estimate was based on “expert knowledge” of the model. To some extent, this takes into account the dataset. However, expert knowledge only takes the amount and type of observed data into account approximately. An alternative, which we recommend, would be to formally consider the choice of parameters to estimate as a problem of model selection, where the selection is of the subset of parameters to estimate by calibration, while the other parameters retain their default values. For example, one could use the Akaike Information Criterion (AIC; Akaike (1973)), which has been widely used for model choice in ecology (Burnham et al., 2011) to choose the parameters to estimate. The use of a model selection rule would automatically adapt the choice of parameters to estimate to the calibration dataset. Consider, for example, the question of whether or not to estimate parameters related to water stress for those models that include effects of water stress on phenology. A model selection rule would include such a parameter if it had a relatively large effect on improving the fit to the calibration data, and would not include it otherwise. However, given the large number of possible parameters to estimate, it would probably be necessary to combine expert knowledge, in order to choose a fairly small number of candidate parameters, with a formal model selection criterion.

All parameters that are not estimated using the calibration data retain their default values, and this, in general, concerns the majority of model parameters. While some parameters will not have an effect on the simulated values, many others will have an effect. It is clear, then, that the choice of these default values is extremely important, and should reflect whatever information one has about the cultivars and environments of interest. The choice of default values probably merits more attention than it usually receives.

### 4.3 Algorithmes and software

There are several disadvantages to the trial and error approach which was used by somewhat over a third of groups. it is time-consuming, it is likely to end in a non-optimal solution, especially if several parameters are estimated, and it cannot be replicated for example to estimate prediction error using cross-validation.

A wide range of algorithms and software was used by the remaining groups. The problem of choosing a calibration algorithm and software to search for optimal parameter values has received much attention in the field of hydrological modeling (Skahill and Doherty, 2006). Gradient based algorithms are, in general, very efficient, but may converge to a local rather than global optimum (Blasone et al., 2006). Also, simulated values are often non-continuous functions of the parameters. As a result, it may not be possible to calculate a gradient. Removing the discontinuities may be possible, but at the price of detailed intervention in the model code (Liu et al., 2018). Global search algorithms, such as a grid search or genetic algorithms, may avoid converging to a local optimum but in general require many more model runs. A third possibility is a gradient-free search algorithm such as the simplex method (Nelder and Mead, 1965), which seems to be a good choice for calibration of crop models.

There is calibration software that has been developed specifically for some crop models (Buddhaboon et al., 2018; Buis et al., 2015; Hunt et al., 1993), and also some software that is designed to be easily coupled to any model (Doherty et al., 2010). Coupling parameter estimation software to a crop model is not simple and so modeling groups tend to use available software or even no software rather than developing new calibration software themselves. This implies that for the improvement of calibration approaches for crop models it is not sufficient to propose guidelines for good calibration practices. For the guidelines to be effective, they must include software solutions that can be used by any model.

### 4.4 Summary of recommendations

Based on an analysis of calibration approach by multiple modeling groups, we propose the following guidelines for calibration of the phenology component of crop models. Before calibration, one should pay careful attention to the choice of default values for all parameters. If the objective is, as here, to make point predictions, then a frequentist approach to calibration is fully justified. One can use an OLS criterion initially, based on errors in units of days, but the statistical assumptions underlying OLS should be checked by analyzing residuals after calibration. The choice of parameters to estimate should be adapted to the available data, for example using the AIC criterion. However, the choice of parameters also requires knowledge of the model and the environments studied, as well as of the reliability of the data. This suggests that a fully automatic calibration procedure may not be advisable. For the calculations, a derivative free but efficient search algorithm like the simplex is recommended. These recommendations are specific to calibration of the phenology component of crop models, but many of them are no doubt more widely applicable.

## 5 Conclusions

Calibration of crop models involves multiple decisions, which can be grouped into choice of criteria for defining the best parameter values, choice of parameters to estimate and choice of algorithm and software. Different modeling groups make quite different decisions, even for modeling groups using the same model structure. It seems that we are far from having a consensus on how to calibrate crop models, even in the relatively simple case with only phenology data, which emphasizes the need for calibration guidelines such as those suggested here.

In developing recommendations for calibration in other fields, we suggest that a procedure similar to ours could be of interest. This consists of obtaining detailed information about calibration procedure from multiple groups, for a specific calibration problem, covering as far as possible the full set of calibration decisions. The recommendations are then based on analyzing the advantages and disadvantages of the different choices that are made. In this way, one obtains recommendations that cover a wide range of the decisions that calibration requires, and that are anchored in actual practice.

## 6 Acknowledgements

This work was in part supported by the Collaborative Research Center 1253 CAMPOS (Project 7: Stochastic Modelling Framework), funded by the German Research Foundation (DFG, Grant Agreement SFB 1253/1 2017), the Academy of Finland through projects AICropPro (316172) and DivCSA (316215) and Natural Resources Institute Finland (Luke) through a strategic project BoostIA, the BonaRes project “Soil3” (BOMA 03037514) of the Federal Ministry of Education and Research (BMBF), Germany, the Deutsche Forschungsgemeinschaft (DFG, German Research Foundation) under Germany’s Excellence Strategy - EXC 2070 – 390732324 (PhenoRob), the project BiomassWeb of the GlobeE programme (Grant number: FKZ031A258B) funded by the Federal Ministry of Education and Research (BMBF, Germany), the INRA ACCAF meta-programme, the German Federal Ministry of Education and Research (BMBF) in the framework of the funding measure “Soil as a Sustainable Resource for the Bioeconomy – BonaRes”, project “BonaRes (Module B): BonaRes Centre for Soil Research, subproject B” (grant 031B0511B), the National Key Research and Development Program of China (2017YFD0300205), the National Science Foundation for Distinguished Young Scholars (31725020), the Priority Academic Program Development of Jiangsu Higher Education Institutions (PAPD), the 111 project (B16026), and China Scholarship Council, the Agriculture and Agri-Food Canada’s Project 1387 under he Canadian Agricultural Partnership, the DFG Research Unit FOR 1695 ‘Agricultural Landscapes under Global Climate Change – Processes and Feedbacks on a Regional Scale, the U.S. Department of Agriculture National Institute of Food and Agriculture (award no. 2015-68007-23133) and USDA/NIFA HATCH grant N. MCL02368, the National Key Research and Development Program of China (2016YFD0300105), The Broadacre Agriculture Initiative, a research partnership between University of Southern Queensland and the Queensland Department of Agriculture and Fisheries, the Academy of Finland through project AI-CropPro (315896), the JPI FACCE MACSUR2 project, funded by the Italian Ministry for Agricultural, Food, and Forestry Policies (D.M. 24064/7303/15 of 6/Nov/2015). The order in which the donors are listed is arbitrary.

## Notes

### Competing Interest Statement

The authors have declared no competing interest.

