## Supplementary material for "The chaos in calibrating crop models": SI

**Table S1**

**Model structures used by participating modeling groups, with references.**

| Model structure | Version(s) | References |
|---|---|---|
| AgroC | May2018 | Herbst M., Hellebrand H.J., Bauer J., Huisman J.A., Šimůnek J., Weihermüller L., Graf A., Vanderborght J., Vereecken H. (2008). Multiyear heterotrophic soil respiration: Evaluation of a coupled $CO_2$ transport and carbon turnover model. Ecological Modelling 214: 271-283.<br>Klosterhalfen A., Herbst M., Weihermüller L., Graf A., Schmidt M., Stadler A., Schneider K., Subke J.-A., Huisman J.A., Vereecken H. (2017). Multi-site calibration and validation of a net ecosystem carbon exchange model for croplands. Ecological Modelling 363: 137-156. |
| APSIM | 7.8, 7.9, 7.10 | Keating B.A., Carberry P.S., Hammer G.L., Probert M.E., Robertson M.J., Holzworth D., Huth N.I., Hargreaves J.N.G., Meinke H., Hochman Z., McLean G., Verburg K., Snow V., Dimes J.P., Silburn M., Wang E., Brown S., Bristow K.L., Asseng S., Chapman S., McCown R.L., Freebairn D.M., Smith C.J. (2003). An overview of APSIM, a model designed for farming systems simulation. European Journal of Agronomy 18: 267-288.<br>Holzworth D.P., Huth N.I., DeVoil P.G. et al. (2014) APSIM - Evolution towards a new generation of agricultural systems simulation. Environmental Modelling & Software 62: 327-350. |
| AquaCrop | 4.0 | Vanuytrecht E., Raes D., Steduto P., Hsiao T.C., Fereres E., Heng L.K., Garcia Vila M., Mejias Moreno P. (2014). AquaCrop: FAO's crop water productivity and yield response model. Environmental Modelling & Software 62: 351-360. |
| CERES-Wheat | DSSAT V4.7, | Hoogenboom G., Porter C.H., Boote K.J., Shelia V., Wilkens P.W., Singh U., White J.W., |

| | | |
|---|---|---|
| | V4.7 | Asseng S., Lizaso J.I., Moreno L.P., Pavan W., Ogoshi R., Hunt L.A., Tsuji G.Y., Jones J.W. (2019). The DSSAT crop modeling ecosystem. In: Boote K.J. (ed.). Advances in Crop Modeling for a Sustainable Agriculture. Burleigh Dodds Science Publishing, Cambridge, United Kingdom, p. 173-216, http://dx.doi.org/10.19103/AS.2019.0061.10.<br>Hoogenboom, G., Porter C.H., Shelia V., Boote K.J., Singh U., White J.W., Hunt L.A., Ogoshi R., Lizaso J.I., Koo J., Asseng S., Singels A., Moreno L.P., Jones J.W. (2019). Decision Support System for Agrotechnology Transfer (DSSAT) Version 4.7 (www.DSSAT.net). DSSAT Foundation, Gainesville, Florida, USA. |
| CoupModel | Version 5.4.4 | Coucheney E., Eckersten H., Hoffmann H., Jansson P.-E., Gaiser T., Ewert F., Lewan E. (2018). Key functional soil types explain data aggregation effects on simulated yield, soil carbon, drainage and nitrogen leaching at a regional scale. Geoderma 318: 167-181. DOI: 10.1016/j.geoderma.2017.11.025.<br>Jansson P.-E. (2012). CoupModel: model use, calibration, and validation. Transactions of the ASABE 55 (4):1337-1344. (American Society of Agricultural and Biological Engineers).<br>Senapati N., Jansson P.-E., Smith P., Chabbi A. (2016). Modelling heat, water and carbon fluxes in mown grassland under multi-objective and multi-criteria constraints. Environmental modelling & software 80: 201-224. |
| CROPSIM-Wheat | DSSAT V4.7 | Hoogenboom, G., Porter C.H., Boote K.J., Shelia V., Wilkens P.W., Singh U., White J.W., Asseng S., Lizaso J.I., Moreno L.P., Pavan W., Ogoshi R., Hunt L.A., Tsuji G.Y., Jones J.W. (2019). The DSSAT crop modeling ecosystem. In: Boote K.J. (ed.). Advances in Crop Modeling for a Sustainable Agriculture. Burleigh Dodds Science Publishing, Cambridge, United Kingdom, p. 173-216, (http://dx.doi.org/10.19103/AS.2019.0061.10.<br>Hoogenboom G., Porter C.H., Shelia V., Boote K.J., Singh U., White J.W., Hunt L.A., Ogoshi R., Lizaso J.I., Koo J., Asseng S., Singels A., Moreno L.P., Jones J.W. (2017). Decision Support System For Agrotechnology Transfer (DSSAT). Version 4.7. DSSAT Foundation, Gainesville, Florida, USA. |
| Cropsyst | 3.04.08 | Stöckle C.O., Donatelli M., Nelson R. (2003). CropSyst, a cropping systems simulation |

| | | |
|---|---|---|
| | | model. European Journal of Agronomy 18(3-4): 289-307. |
| DAISY | 5.59 | Hansen S., Abrahamsen P., Petersen C.T., Styczen M. (2012). Daisy: Model Use, Calibration, and Validation. Transactions of the ASABE 55: 1317-1335. |
| Nwheat | DSSAT | Kassie B.T., Asseng S., Porter C.H. and Royce F.S. (2016). Performance of DSSAT-Nwheat across a wide range of current and future growing conditions. European Journal of Agronomy 81: 27-36.<br>Hoogenboom, G., Porter C.H., Boote K.J., Shelia V., Wilkens P.W., Singh U., White J.W., Asseng S., Lizaso J.I., Moreno L.P., Pavan W., Ogoshi R., Hunt L.A., Tsuji G.Y., Jones J.W. (2019). The DSSAT crop modeling ecosystem. In: Boote K.J. (ed.). Advances in Crop Modeling for a Sustainable Agriculture. Burleigh Dodds Science Publishing, Cambridge, United Kingdom, p. 173-216, http://dx.doi.org/10.19103/AS.2019.0061.10. |
| GECROS | Expert-N 3.0 | Yin X., van Laar H.H. (2005). Crop systems dynamics. An ecophysiological simulation model for genotype-by-environment interactions. Wageningen Academic Publishers, 155 pp., Wageningen, The Netherlands. |
| HERMES | 4.27 | Kersebaum K.C. (2007). Modelling nitrogen dynamics in soil-crop systems with HERMES. Nutrient Cycling in Agroecosystems 77: 39-52.<br>Kersebaum K.C. (2011). Special features of the HERMES model and additional procedures for parameterization, calibration, validation, and applications In: Ahuja L.R., Ma L. (eds.): Advances in Agricultural Systems Modeling Series 2. 65-94. ASA, CSSA, SSSA, Madison, USA. |
| LINTUL | LINTUL5 | Wolf J. (2012). User guide for LINTUL5: Simple generic model for simulation of crop growth under potential, water limited and nitrogen, phosphorus and potassium limited conditions. Wageningen UR. |
| MONICA | 2.02 | Nendel C., Berg M., Kersebaum K.C., Mirschel W., Specka X., Wegehenkel M., Wenkel K.O., |

| | | |
|---|---|---|
| | | Wieland R. (2011). The MONICA model: Testing predictability for crop growth, soil moisture and nitrogen dynamics. Ecological Modelling 222(9): 1614-1625.<br>Specka X., Nendel C., Wieland R. (2015). Analysing the parameter sensitivity of the agro-ecosystem model MONICA for different crops. European Journal of Agronomy 71: 73-87.<br>Specka X., Nendel C., Wieland R. (2019). Temporal sensitivity analysis of the MONICA Model: application of two global approaches to analyze the dynamics of parameter sensitivity. Agriculture 9(2): 37. |
| OpenCrop | | Crout N.M.J., Karanaratne A, Jabloun M. (2018). OpenCrop: An Open Source Crop Model – Model Description. School of Biosciences, University of Nottingham, UK. |
| PANORAMIX | R version | Gate P. (1995). Écophysiologie du blé. Lavoisier-Technique et documentation.<br>Chatelin M.H., Aubry C., Poussin J.C., Meynard J.M., Massé J., Verjux N., Gate P., Le Bris X., 2005. DéciBlé, a software package for wheat crop management simulation. Agricultural Systems 83: 77-99, https://doi.org/10.1016/J.AGSY.2004.03.003. |
| Salus | | Basso B., Ritchie J.T., Grace P.R., Sartori L. (2006). Simulation of tillage systems impact on soil biophysical properties using the SALUS model. Italian Journal of Agronomy 1: 677-688.<br>Basso B., Ritchie J.T. (2015). Simulating crop growth and biogeochemical fluxes in response to land management using the SALUS Model. In: Hamilton S.K., Doll J.E., Robertson G.P. (eds). The ecology of agricultural landscapes: long-term research on the path to sustainability. Oxford University Press, New York, NY USA. |
| SPASS | Expert-N 3.0 | Wang E. (1997). Development of a Generic Process-Oriented Model for Simulation of Crop Growth. München, Herbert Utz Verlag Wissenschaft. 195 pp. |
| SSM-Wheat | | Soltani A., Maddah V., Sinclair T. (2013). SSM-Wheat: a simulation model for wheat development, growth and yield. International Journal of Plant Production 7: 711-740. |
| STICS | 8_5_0 | Brisson N., Launay M., Mary B., Beaudoin N. (2009). Conceptual basis, formalisations and |

| | | |
|---|---|---|
| | | parameterization of the STICS crop model. Quae, 304pp.<br>Coucheney E., Buis S., Launay M., Constantin J., Mary B., Garcia de Cortazar-Atauri I., Ripoche D., Beaudoin N., Ruget F., Andrianorisoa S., Le Bas C., Justes E., Léonard J. (2015). Accuracy, robustness and behavior of the STICS 8.2.2 soil-crop model for plant, water and nitrogen outputs: evaluation over a wide range of agro-environmental conditions in France. Environmental Modelling & Software 64: 177-190. |
| SUCROS | Expert-N 3.0 | van Laar H.H., Goudriaan J., van Keulen H. (1992). Simulation of crop growth for potential and water-limited production situations (as applied to spring wheat). Simulation Report CABO-TT no. 27. Wageningen: Centre for Agrobiological Research and Department of Theoretical Production Ecology, Wageningen Agricultural University.<br>Vanclooster M., Viaene P., Diels J., Christiaens K. (1994). WAVE a mathematical model for simulating water and agrochemicals in the soil and vadose environment. Reference and user's manual (release 2.0). Leuven: Institute for Land and Water Management, Katholieke Universiteit Leuven. |
| PCWOFOST | 5.3.3 | Ceglar A., van der Wijngaart R., de Wit A., Lecerf R., Boogaard H., Seguini L., van den Berg M., Toreti A., Zampieri M., Fumagalli D., Baruth B. (2019). Improving WOFOST model to simulate winter wheat phenology in Europe: Evaluation and effects on yield. Agricultural Systems 168: 168-180. |
| WCCWOFOST | 7.1.7 | Boogaard H.L., Van Diepen C.A., Rötter R.P., Cabrera J.M.C.A., Van Laar H.H. (1998). User's guide for the WOFOST 7.1 crop growth simulation model and WOFOST control center 1.5. Technical Document 52. Winand Staring Centre, Wageningen, the Netherlands, 144 pp. |
| Wheat-Grow | 3.1 | Zhu Y., Liu L., Liu B. (2013). WheatGrow: A simulation model for predicting growth and productivity in wheat. In: Proceedings of the Workshop on Modeling Wheat Response to High Temperature, Texcoco, Mexico, 19–21 June 2013.<br>Lv Z., Liu X., Tang L., Liu L., Cao W., Zhu, Y. (2016). Estimation of ecotype-specific cultivar parameters in a wheat phenology model and uncertainty analysis. Agricultural and Forest |

| | | |
|---|---|---|
| | | Meteorology 221: 219-229. |

.

**Table S2**

**The questionnaire filled out by all participating groups concerning their calibration procedure**

### Explanation of calibration procedure

Please answer the following questions. The objective is to better understand exactly how calibration was done. We are particularly interested in 5 aspects of calibration: choice of parameters (which, how many), fixing lower and upper limits on parameter values or other prior information, how multiple measured variables were handled, the calibration approach (e.g. weighted least squares, Bayes, GLUE etc), calculation of uncertainty in estimated parameters.

1. Name of model. Version number.
2. Contact person, name and email
3. How many parameters did you estimate (i.e. change from initial default values)?
4. Which parameters did you estimate (please give the name and a short explanation and units of each)?
5. Why did you choose those particular parameters to calibrate? Why that number of parameters to estimate (i.e. why not more or fewer parameters)?
6. Which of the measured variables did you use for calibration (e.g. both date of stem elongation and of heading)?
7. Describe your calibration approach. For example, did you define a criterion to minimize, like $\sum_{i=1}^{14}\left[\left(y_{i,BBCH30}^{obs}-y_{i,BBCH30}^{sim}\right)^2+\left(y_{i,BBCH55}^{obs}-y_{i,BBCH55}^{sim}\right)^2\right]$ i.e. the total sum of squared errors for days to BBCH30 and BBCH55, where 14 is the number of training environments . The variables specified in response to question 6, and no others, should appear here.
8. Did you set lower and upper limits to the parameter values, or use other prior information about the parameter values? If so, please give the values for each parameter. How did you decide on those limits or that prior distribution?
9. What software did you use to find the values of the parameters?
10. Did you estimate parameter uncertainty? If so, please give the uncertainty values for your parameters (for example, standard deviation for each parameter), and indicate how you estimated the uncertainty.

11. We provided as input variables multiple variables for weather (daily max and min T etc), multiple variables for soil characteristics (e.g. water holding capacity by layer), management (sowing date, irrigation, fertilization) and initial conditions (water, NO3). Did your model require other input variables that weren't provided with the data? Please list them and give the values you assigned them. (We are interested to see if different groups make different decisions here, which could help explain why different groups running the same model obtain different results). Also, please indicate if you made changes to the input data we provided, and explain why.
12. Which of the following affect phenology according to the model you used:

Daily average temperature

Hourly temperature

Heat stress (duration of maximum temperature above some threshold)

Vernalization

Day length

Solar radiation

Water stress

N stress

Other (please specify)

13. Any other comments

#### Table S3

**Details of choices by each modeling group concerning the criteria for defining the best parameters, for the French and Australian datasets.**

**[1, 2, 3] Groups with the same superscript used the same model structure, namely S1, S2, or S3, respectively**

a. **The variables to be predicted are BBCH30 and BBCH55**
b. **Errors can be measured in days or in development stage. The choice was days unless explicitly stated otherwise.**
c. **The variables to be predicted are Z30, Z65 and Z90**
d. **This group assumed a constant thermal time from Z55 to Z65, so this is equivalent to using observations of days from sowing to Z30 and to Z65.**

| Modeling group | French dataset | | Australian dataset | |
|---|---|---|---|---|
| | Variables included in criterion[a]<br><br>Units of error[b] | Frequentist or Bayes<br><br>Objective function or Likelihood and priors | Variables included in criterion[c]<br><br>Units of error[b] | Frequentist or Bayes<br><br>Objective function or Likelihood and priors |
| M1 | BBCH30, BCH55 | Frequentist<br><br>Sum of squared errors | Z13, Z16, Z19, Z22, Z27, Z31, Z36, Z42, Z48, Z54, Z60, Z66, Z72, Z78, Z83, Z90, Z98 | Frequentist<br><br>Sum of squared errors |
| M2[1] | BBCH30, BCH55 | Frequentist<br><br>Sum of squared errors | Z15, Z30, Z40, Z50, Z60, Z65, Z70, Z80, Z90 | Frequentist<br><br>Sum of squared errors |

| | | | | |
|---|---|---|---|---|
| M3[1] | BBCH30, BCH55 | Frequentist<br><br>Sum of squared errors | Z65, Z90 | Frequentist<br><br>Sum of squared errors |
| M4[1] | BBCH30, BCH55 | Frequentist<br><br>Sum of squared errors | Z30, Z65, Z90 | Frequentist<br><br>Sum of squared errors |
| M5[1] | BBCH30, BCH55 | Frequentist<br><br>Sum of squared errors | Didn't participate | |
| M6 | BBCH30, BCH55 | Frequentist<br><br>Sum of squared errors | Z30, Z65, Z90 | Frequentist<br><br>Sum of squared errors |
| M7[2] | BBCH30, BCH55 | Frequentist<br><br>Sum of squared errors<br><br>Sequential fit: BBCH30 then BBCH55 | All data | Frequentist<br><br>Sum of squared errors<br><br>Sequential fit: Z30 then Z65, then Z90, then others |
| M8 | BBCH30, BCH55 | Frequentist<br><br>Sum of squared errors<br><br>Sequential fit: BBCH30, then BBCH55 | All data | Frequentist<br><br>Sum of squared errors |
| M9 | BCH55 | Frequentist<br><br>Sum of squared errors | Z65, Z71, Z90 | Frequentist<br><br>Sum of squared errors<br><br>Sequential fit: each stage treated separately |

| | | | | |
|---|---|---|---|---|
| M10 | BBCH30, BCH55 | Frequentist<br>Sum of squared errors | Z65, Z90 | Frequentist<br>Sum of squared errors |
| M11 | BBCH30, BCH55 | Frequentist<br>Sum of squared errors | Z30, Z65, Z90 | Frequentist<br>Sum of squared errors |
| M12[2] | BBCH30, BCH55 | Frequentist<br>No single objective<br>First fit each environment separately, then average parameters over environments | All data | Frequentist<br>No single objective<br>First fit each environment separately, then average parameters over environments |
| M13[2] | BBCH30, BCH55 | Frequentist<br>No single objective | Z30, Z65 | Frequentist<br>No single objective |
| M14 | BBCH30, BCH55 | Frequentist<br>Sum of squared errors | First available Zadoks stage, last available Zadoks stage, Z65<br>Error in units of development stage | Frequentist<br>Sum of squared errors |
| M15 | BBCH30, BCH55 | Frequentist<br>Sum of root mean squared errors | Z30, Z55, Z65, Z90 | Frequentist<br>Sum of root mean squared errors |
| M16 | BBCH30, BCH55 | Frequentist<br>No single objective | Z65, Z90 | Frequentist<br>No single objective |

| | | | | |
|---|---|---|---|---|
| M17 | BBCH30, BCH55 | Frequentist<br><br>Sum of squared errors | Z12, Z30, Z59, Z60, Z65, Z69, Z70, Z89, Z90 | Frequentist<br><br>Sum of squared errors |
| M18 | BCH55 | Frequentist<br><br>Sum of squared errors | Z65, Z90 | Frequentist<br><br>Sum of root mean squared errors |
| M19 | BBCH30, BCH55 | Bayesian<br><br>Normal likelihood; iid errors.<br><br>Uniform priors | Z30, Z65, Z90 | Bayesian<br><br>Normal likelihood.<br><br>Assume standard deviation of model error for each variable is proportional to 1/(number of observations)<br><br>Uniform priors |
| M20 | BBCH30, BCH55 | Frequentist<br><br>No single objective<br><br>(Look at normalized root mean squared error, mean absolute error, and $R^2$) | Z21, Z31, Z40, Z55, Z65, Z70, Z80, Z90 | Frequentist<br><br>No single objective<br><br>(Look at bias, root mean squared error, $R^2$, and slope of observed vs simulated values) |
| M21 | BBCH30, BCH55 | Bayesian<br><br>Concentrated likelihood<br><br>Unform or truncated normal priors | Z30, Z65, Z71, Z90 | Bayesian<br><br>Concentrated likelihood<br><br>Uniform priors |

| | | | | |
|---|---|---|---|---|
| M22 | BBCH30, BCH55 | Frequentist<br><br>Sum of squared errors | Z30, Z55, Z65, Z90 | Frequentist<br><br>Sum of squared errors |
| M23[3] | BBCH30, BCH55 | Frequentist<br><br>Sum of squared errors | Z30, Z65, Z90 | Frequentist<br><br>Sum of squared errors |
| M24[3] | BBCH30, BCH55 | Frequentist<br><br>Sum of squared errors | Z30, Z65, Z90 | Frequentist<br><br>Sum of squared errors<br><br>Sequential fit: Z30 and Z65 together, then Z90 |
| M25[2] | BBCH30, BCH55 | Frequentist<br><br>Sum of squared errors | All data<br><br>(one environment couldn't be fitted, so wasn't used)<br><br>Error in units of development stage | Bayesian<br><br>Likelihood has Student's *t* distribution<br><br>Uniform priors |
| M26 | BBCH30, BCH55 | Frequentist<br><br>Sum of squared errors | All data | Frequentist<br><br>Weighted sum of squared errors<br><br>weight=10 for Z30, Z65, Z90<br><br>weight=6 for first, last provided development stage<br><br>weight=1 otherwise |
| M27 | BBCH30, BCH55 | Frequentist | All data | Frequentist |

| | | | | |
|---|---|---|---|---|
| | | Sum of squared errors | | Weighted sum of squared errors<br>weight=10 for Z30, Z65, Z90<br>weight=6 for first, last provided development stage<br>weight=1 otherwise |
| M28 | Didn't participate | | All data<br>Error in units of development stage | Bayesian<br>Normal likelihood<br>Normal priors |
| M29 | Didn't participate | | Z30, Z55[d] | Frequentist<br>Sum of squared errors<br>Sequential fit: Z30 and then difference Z65-Z30 |

#### Table S4

**Details of choices by each modeling group concerning the selection of parameters to estimate, for the French and Australian datasets.**

**[1, 2, 3] Groups with the same superscript used the same model structure, namely S1, S2, or S3, respectively.**

**a. Number of estimated parameters**

**b. PT=physiological time, which is a measure of rate of development and usually depends on temperature, possibly modified by other factors such as photoperiod, from sowing up to the indicated development stage or between the indicated development stages, photoperiod=parameter(s) related to response of development rate to photoperiod, vernalization=parameter(s) related to accumulation of cold temperatures required for flowering, base temperature=minimum temperature for development, optimal temperature=optimal temperature for development.**

| | French dataset | | | Australian dataset | | |
|---|---|---|---|---|---|---|
| Modeling group | Basis for choosing parameters to estimate | N[a] | Estimated parameters[b] | Basis for choosing parameters to estimate | N[a] | Estimated parameters[b] |
| M1 | Expert knowledge and data based (adjusted $R^2$, AIC) | 1 | Vernalization | Expert knowledge | 3 | Temperature scaling factor DVS<1<br>Temperature scaling factor DVS>1<br>Vernalization |

| | | | | | | |
|---|---|---|---|---|---|---|
| M2[1] | Expert knowledge | 3 | PT floral initiation to flowering<br>Vernalization<br>Photoperiod | Expert knowledge | 5 | PT floral initiation to flowering<br>PT start to end flowering<br>PT start to end grain fill<br>Vernalization<br>Photoperiod |
| M3[1] | Expert knowledge | 2 | Vernalization<br>photoperiod | Expert knowledge | 2 | Vernalization<br>Photoperiod |
| M4[1] | Expert knowledge | 2 | Vernalization<br>Photoperiod | Expert knowledge | 7 | PT floral initiation<br>PT flower<br>PT start grain fill<br>PT end grain fill<br>PT start grain fill to maturity<br>Vernalization<br>Photoperiod |
| M5 | Expert knowledge | 9 | Thermal time end juvenile to floral initiation<br>Thermal time flowering to start grain fill<br>Thermal time start to end grain fill<br>Thermal time start grain fill to maturity<br>Vernalization<br>Photoperiod<br>Phyllochron<br>Kernel number/stem weight<br>Potential kernel growth rate | Didn't participate | | |
| M6 | Sensitivity analysis | 6 | PT sowing to maturity<br>PT sowing to emergence<br>PT sowing to flowering<br>Model development stage at BBCH30<br>Model development sage at BBCH55<br>Tmax for development rate | Sensitivity analysis | 8 | PT sowing to maturity<br>PT sowing to emergence<br>PT sowing to flowering<br>Tmax for development rate<br>Soil water threshold<br>Crop coefficient for closed canopy |

| | | | | | | |
|---|---|---|---|---|---|---|
| | | | | | | Minimum rooting depth |
| M7[2] | Expert knowledge and data-based (test various choices) | 4 | PT end juvenile to terminal spike<br>Leaf number at start tillering<br>Tillering rate<br>Phyllochron | Expert knowledge and data-based (test various choices) | 9 | PT end juvenile to terminal spikelet<br>PT end leaf growth to end spike growth<br>PT end spike growth to anthesis<br>PT start to end anthesis<br>PT end spike growth to end grain fill lag<br>PT grain filling<br>Vernalization<br>Leaf number at start tillering<br>Tillering rate<br>Phyllochron |
| M8 | Expert knowledge and data-based (test various choices) | 4 | PT end juvenile to terminal spikelet<br>Leaf number at start tillering<br>Tillering rate<br>Phyllochron | Expert knowledge and data-based (test various choices) | 8 | PT end juvenile to terminal spikelet<br>PT end leaf growth to end spike growth<br>PT flowering duration<br>PT grain filling<br>Vernalization<br>Leaf number at start tillering<br>Tillering rate<br>phyllochron |
| M9 | Expert knowledge | 3 | PT to anthesis<br>Vernalization<br>Effect water stress | Expert knowledge | 3 | PT to flowering<br>PT to start grain filling<br>PT to maturity |
| M10 | Sensitivity analysis | 3 | Development rate vegetative phase<br>Development rate reproductive phase<br>Soil temperature sum for emergence | Sensitivity analysis | 3 | Development rate vegetative phase<br>Development rate reproductive phase<br>Soil temperature sum for emergence |
| M11 | Expert knowledge | 3 | PT emergence to end juvenile<br>Vernalization | Expert knowledge | 3 | PT emergence to end juvenile<br>Vernalization |

| | | | | | | |
|---|---|---|---|---|---|---|
| | | | Photoperiod | | | Photoperiod |
| M12[2] | Expert knowledge (Follow guidelines for this model structure) | 3 | PT linear grain filling to maturity<br>Vernalization<br>Photoperiod | Expert knowledge (Follow guidelines for this model structure) | 4 | PT end juvenile to terminal spikelet<br>PT linear grain filling to maturity<br>Vernalization<br>Photoperiod |
| M13[2] | Expert knowledge | 2 | PT end juvenile to terminal spikelet<br>PT end leaf growth to end spike growth | Expert knowledge | 2 | PT end juvenile to terminal spikelet<br>PT end leaf growth to end spike growth |
| M14 | Sensitivity analysis | 6 or 7 | PT vegetative phase<br>Photoperiod<br>Development stage at start of photoperiod sensitivity (Bermude)<br>Development stage at end of photoperiod sensitivity (Bermude)<br>Base temperature (Apache)<br>Optimal temperature<br>Ceiling temperature<br>Coefficient of temperature response | Sensitivity analysis | 4 | PT vegetative phase<br>PT reproductive phase<br>Photoperiod<br>Optimal temperature |
| M15 | Expert knowledge | 5 | PT sowing to emergence<br>PT emergence to double ridge<br>Vernalization<br>Photoperiod for phase 2<br>Photoperiod for phase 3 | Expert knowledge | 10 | PT sowing to emergence<br>PT emergence to double ridge<br>PT double ridge to heading<br>PT heading to flowering<br>PT flowering to maturity<br>Vernalization<br>Photoperiod phase 2<br>Photoperiod phase 3<br>Base temperature phase 1 |

| | | | | | | |
|---|---|---|---|---|---|---|
| | | | | | | Base temperature phase 4 |
| M16 | Expert knowledge | 2 | PT sowing to emergence<br>Vernalization | Expert knowledge | 4 | PTsowing to emergence<br>PT emergence to anthesis<br>PT anthesis to maturity<br>Vernalization |
| M17 | Expert knowledge | 2 | PT emergence to double ridge<br>PT double ridge to heading | Expert knowledge | 5 | PT emergence to double ridge<br>PT double ridge to heading<br>PT double ridge to flowering<br>PT flowering to grain filling<br>PT grain filling |
| M18 | Expert knowledge | 2 | PT sowing to flowering<br>Base temperature | Expert knowledge | 2 | PT sowing to flowering<br>PT sowing to maturity mat |
| M19 | Sensitivity analysis | 3 | Photoperiod<br>Phyllochron<br>Cold tolerance | Sensitivity analysis | 2 | Photoperiod<br>Phyllochron |
| M20 | Expert knowledge | 6 | PT tillering to stem elongation<br>PT stem elongation to booting<br>PT booting to ear emergence<br>Vernalization<br>Photoperiod<br>Phyllochron | Expert knowledge | 8 | PT emergence to tillering<br>PT tillering to stem elongation<br>PT stem elongation to booting<br>PT booting to heading<br>PT heading to anthesis<br>PT anthesis to maturity<br>Vernalization<br>Photoperiod |
| M21 | Expert knowledge | 3 | PT emergence to end juvenile<br>PT end juvenile to maximum LAI<br>Vernalization | Expert knowledge | 6 | PT emergence to end juvenile<br>PT emergence to start grain filling<br>PT anthesis to start grain filling<br>PT start grain filling to maturity<br>Vernalization<br>Maximum delay due to stresses |

| | | | | | | |
|---|---|---|---|---|---|---|
| M22 | Expert knowledge | 4 | Intrinsic earliness<br>Vernalization<br>Photoperiod<br>Thermal sensitivity | Expert knowledge | 5 | Intrinsic earliness<br>Filling duration<br>Vernalization<br>Photoperiod<br>Thermal sensitivity |
| M23[3] | Expert knowledge | 3 | PT emergence to flowering<br>Model development stage at BBCH30<br>Model development stage at BBCH65 | Expert knowledge | 4 | PT sowing to emergence<br>PT emergence to flowering<br>PT flowering to maturity<br>Model development stage at Z30 |
| M24[3] | Expert knowledge and data-based (test various choices) | 1 | PT emergence to flowering | Expert knowledge and data-based (test various choices) | 2 | PT emergence to flowering<br>PT flowering to maturity |
| M25[2] | Expert knowledge | 5 | PT stage 1<br>PT stage 4<br>Vernalization<br>Photoperiod<br>Phyllochron | Expert knowledge | 8 | PT stage 1<br>PT stage 4<br>PT stage 5<br>Vernalization<br>Photoperiod<br>Phyllochron<br>Standard deviation of residuals<br>Degrees of freedom of t-distribution |
| M26 | Sensitivity analysis | 6 | PT vegetative phase<br>Vernalization<br>Photoperiod sensitivity<br>Photoperiod optimal<br>Optimum temperature<br>Base temperature | Expert knowledge | 7 | PT vegetative<br>PT reproductive<br>Vernalization<br>Photoperiod<br>Optimal temperature during vegetative phase<br>Base temperature during reproductive |

| | | | | | | |
|---|---|---|---|---|---|---|
| | | | | | | phase |
| M27 | Expert knowledge | 2 | PT emergence anthesis<br>Tbase | Expert knowledge | 4 | PT emergence anthesis<br>PT anthesis mat<br>Base temperature during vegetative phase<br>Base temperature during reproductive phase |
| M28 | Didn't participate | - | - | Expert knowledge | 7 | PT emergence<br>PT emergence to start grain filling<br>PT maturity<br>Zadoks stage at model stage 2<br>Photoperiod<br>Coefficient of temperature response<br>Soil moisture for emergence |
| M29 | Didn't participate | - | - | Expert knowledge and data-based (test various choices) | 5 | PT to Z30<br>PT Z90<br>Three cardinal temperatures up to Z30 |

**Table S5**

**Details of choices made by each modeling group concerning algorithm and software for the French and Australian datasets.**

**[1, 2, 3] Groups with the same superscript used the same model structure, namely S1, S2, or S3, respectively.**

| Modeling group | French data | | Australian data | |
|---|---|---|---|---|
| | Algorithm<br>Software | Bounds on parameter values? | Algorithm<br>Software | Bounds on parameter values? |
| M1 | Global search<br>SCE-UA | Yes | Global search<br>SCE-UA | Yes |
| M2[1] | Gradient based<br>PEST | Yes | Gradient based<br>PEST | Yes |
| M3[1] | Trial and error<br>- | Yes | Trial and error<br>- | Yes |
| M4[1] | Trial and error<br>- | Yes | Trial and error<br>- | Yes |
| M5[1] | Gradient based<br>PEST | Yes | Didn't participate | |
| M6 | MCMC<br>DREAM | Yes | MCMC<br>DREAM | Yes |
| M7[2] | Trial and error<br>- | Yes | Trial and error<br>- | Yes |
| M8 | Trial and error<br>- | Yes | Trial and error<br>- | Yes |
| M9 | Grid search<br>- | No | Grid search<br>- | - |
| M10 | Global search | Yes | Global search | Yes |

| | | | | |
|---|---|---|---|---|
| | DEOptim | | DEOptim | |
| M11 | Trial and error<br>- | Yes | Trial and error<br>- | Yes |
| M12[2] | Trial and error<br>- | Yes | Trial and error<br>- | Yes |
| M13[2] | Trial and error<br>- | No | Trial and error<br>- | No |
| M14 | Gradient based<br>ucode | Yes | Gradient based<br>ucode | Yes |
| M15 | Trial and error<br>- | Yes | Trial and error<br>- | Yes |
| M16 | Trial and error<br>- | - | Trial and error<br>- | No |
| M17 | Global search<br>SCE-UA | - | Global search<br>SCE-UA | Yes |
| M18 | Global search<br>DEOptim | Yes | Global search<br>DEOptim | Yes |
| M19 | MCMC<br>- | Yes | MCMC<br>- | Yes |
| M20 | Trial and error<br>- | No | Trial and error<br>- | Yes |
| M21 | MCMC<br>DREAM | Yes | MCMC<br>DREAM | Yes |
| M22 | Trial and error<br>- | - | Trial and error<br>- | No |
| M23[3] | Global search then gradient based<br>DIRECT-L then gradient method in NLopt library | - | Global search then gradient based<br>DIRECT-L then gradient method in NLopt library | Yes |

| | | | | |
|---|---|---|---|---|
| M24[3] | Trial and error<br>- | Yes | Trial and error<br>- | No |
| M25[2] | Global search<br>DEOptim | Yes | MCMC<br>DRAM | Yes |
| M26 | Gradient-based<br>ucode | Yes | Gradient-based<br>ucode | Yes |
| M27 | Gradient based<br>ucode | Yes | Gradient based<br>ucode | Yes |
| M28 | Didn't participate | | MCMC<br>Software provided with model | Yes |
| M29 | Didn't participate | | Search<br>Nelder-Mead in R function optim | No |

**Table S6**

**Software used for calibration**

| Software name | Description (according to reference) | References |
|---|---|---|
| DEoptim | R Package for Global Optimization by Differential Evolution. The DEoptim function of the package DEoptim searches for minima of the objective function between lower and upper bounds on each parameter to be optimized. DEoptim is particularly well-suited to find the global optimum of a real-valued function of real-valued parameters, and does not require that the function be either continuous or differentiable. | Mullen, K., Ardia, D., Gil, D., Windover, D., Cline, J., 2011. "DEoptim": An R Package for Global Optimization by Differential Evolution. J. Stat. Softw. 40, 1–26. |
| DIRECT-L | DIRECT_L is in the Python NLopt library which has deterministic-search algorithms based on systematic division of the search domain into smaller and smaller hyperrectangles. | Johnson, S.G., The NLopt nonlinear-optimization package, http://github.com/stevengj/nlopt |
| DRAM | The DRAM algorithm combines two quite powerful ideas that have recently appeared in the Markov chain Monte Carlo (MCMC) literature: adaptive Metropolis samplers and delaying rejection. The method is available in the R package FME. | Haario, H., Laine, M., Mira, A., Saksman, E., 2006. DRAM: Effcient Adaptive MCMC. Stat. Comput. 16, 339–354. |
| DREAM | DiffeRential Evolution Adaptive Metropolis (DREAM). This multi-chain MCMC simulation algorithm automatically tunes the scale and orientation of the proposal distribution en route to the target distribution, and exhibits excellent sampling efficiencies on complex, high-dimensional, and multi-modal target distributions. | Vrugt, J.A., ter Braak, C.J.F., Diks, C.G.H., Robinson, B.A., Hyman, J.M., Higdon, D., 2009. Accelerating Markov Chain Monte Carlo Simulation by Differential Evolution with Self-Adaptive Randomized Subspace Sampling. Int. J. Nonlinear Sci. Numer. Simul. 10. https://doi.org/10.1515/IJNSNS.2009.10.3.273 |

| | | |
|---|---|---|
| Nelder-Mead | This is a simplex search algorithm that doesn't require derivatives. | Nelder, J.A., Mead, R., 1965. A Simplex Method for Function Minimization. Comput. J. 7, 308–313. |
| NLOpt library | NLopt includes implementations of a number of different optimization algorithms. | Johnson, S.G., The NLopt nonlinear-optimization package, http://github.com/stevengj/nlopt |
| PEST | An automatic calibration procedure that minimizes an objective function related to the square difference between a number of observed and simulated variables. | Doherty, J.E., Hunt, R.J., Tonkin, M.J., 2010. Approaches to highly parameterized inversion: A guide to using PEST for model-parameter and predictive-uncertainty analysis: U.S. Geological Survey Scientific Investigations Report 010–5211. |
| SCE-UA | The shuffled complex evolution algorithm (SCE-UA) is an MCMC algorithm that reduces the risk of getting stuck in a local optimum by starting several chains/complexes that evolve individually in the parameter space. The population is periodically shuffled and new complexes are created. SCE-UA has been found to be very robust in finding the global optimum of hydrological models. | Houska, T., Kraft, P., Chamorro-Chavez, A., Breuer, L., 2015. SPOTting Model Parameters Using a Ready-Made Python Package. PLoS One 10, e0145180. https://doi.org/10.1371/journal.pone.0145180<br><br>Duan, Q.Y., Gupta, V.K., Sorooshian, S., 1993. Shuffled complex evolution approach for effective and efficient global minimization. J. Optim. Theory Appl. 76, 501–521. https://doi.org/10.1007/BF00939380 |

| | | |
|---|---|---|
| ucode | Minimizes a weighted least-squares objective function with respect to the parameter values using a modified Gauss-Newton method. | Poeter, E.P., Hill, M.C., 1999. UCODE, a computer code for universal inverse modeling. Comput. Geosci. 25, 457–462. https://doi.org/10.1016/S0098-3004(98)00149-6<br><br>Poeter, E.P., Hill, M.C., Banta, E.R., Mehl, S., Christensen, S., 2005. UCODE_2005 and Six Other Computer Codes for Universal Sensitivity Analysis, Calibration, and Uncertainty Evaluation: U.S. Geological Survey Techniques and Methods 6-A11. |

**Table S7**

**Mean absolute error (MAE) in days for each model, separately for the French and Australian datasets and for the calibration and evaluation environments. Emedian, emean, onlyT and naïve represent respectively the added reference models that predict using the median of predictions by individual models, using the mean of predictions by individual models, using a simplified temperature sum model and using the mean of the results for the calibration environments.**

**[1, 2, 3] Groups with the same superscript used the same model structure, namely S1, S2, or S3, respectively.**

| | French dataset | | Australian dataset | |
|---|---|---|---|---|
| | MAE eval | MAE cal | MAE_eval | MAE_cal |
| M1 | 9.00 | 6.23 | 15.8 | 12.8 |
| M2[S1] | 3.94 | 3.80 | 6.8 | 6.3 |
| M3[S1] | 4.91 | 5.36 | 8 | 7.7 |

| | | | | |
|---|---|---|---|---|
| M4[S1] | 6.75 | 6.68 | 6.7 | 5.7 |
| M5[S1] | 12.75 | 17.98 | NA | NA |
| M6 | 12.81 | 10.29 | 14 | 8.1 |
| M7[S2] | 5.97 | 4.71 | 9.3 | 13.3 |
| M8 | 5.53 | 5.09 | 9.4 | 12 |
| M9 | 9.31 | 6.29 | 6.3 | 6.2 |
| M10 | 6.19 | 5.98 | 10.6 | 12 |
| M11 | 9.31 | 11.21 | 7.3 | 9 |
| M12[S2] | 7.03 | 9.34 | 9.3 | 8.1 |
| M13[S2] | 4.31 | 4.73 | 7.2 | 8.3 |
| M14 | 5.06 | 4.68 | 13 | 18.7 |
| M15 | 3.72 | 3.09 | 7.3 | 8 |
| M16 | 6.09 | 5.20 | 14.2 | 13.9 |
| M17 | 3.47 | 3.77 | 8.4 | 7 |
| M18 | 10.50 | 6.11 | 7.2 | 7.7 |
| M19 | 10.69 | 11.77 | 8.5 | 7.9 |
| M20 | 6.41 | 4.43 | 9.3 | 8.3 |
| M21 | 4.81 | 3.43 | 6.6 | 5.9 |
| M22 | 7.78 | 4.88 | 9.5 | 7.3 |
| M23[S3] | 4.66 | 3.52 | 7.8 | 6.7 |
| M24[S3] | 5.00 | 6.20 | 6.4 | 8.5 |
| M25 | 4.78 | 3.70 | 7.4 | 6.1 |
| M26 | 5.00 | 4.20 | 7.9 | 7.4 |
| M27 | 9.59 | 6.30 | 8 | 6.7 |

| | | | | |
|---|---|---|---|---|
| M28 | NA | NA | 20 | 17.4 |
| M29 | NA | NA | 8.2 | 9.7 |
| emedian | 3.53 | 3.68 | 6.4 | 5.9 |
| emean | 4.03 | 3.91 | 6.3 | 6 |
| onlyT | 11.03 | 8.07 | 8.2 | 8 |
| naive | 11.05 | 8.63 | 11.3 | 17.1 |
